# Inhalation Delivered RNA Aptamer for Therapeutics against SARS-CoV2

**DOI:** 10.1101/2023.10.28.564096

**Authors:** Minsun Song, Jiehua Zhou, Alberto Herrera, Peter Baines, John J. Rossi

## Abstract

The continual emergence and re-emergence of infectious diseases has led to a pressing need for the development of swift and targeted therapeutic interventions. SARS-CoV-2, the causative agent of COVID-19, is a prime example of such a rapidly spreading virus[1]. The global crisis caused by this virus has propelled researchers to explore and adopt novel techniques in the hopes of effectively combating its spread[2]. One such innovative approach is the use of aptamers—single-stranded RNA molecules that can bind targets with high specificity.

## Introduction

Aptamers, often dubbed chemical antibodies, are garnered through a process known as Systematic Evolution of Ligands by Exponential Enrichment (SELEX)[3-5]. This method is effective in isolating high-affinity single-stranded RNAs from a vast library of randomized sequences. Like antibodies, aptamers have a keen ability to bind to an array of targets, ranging from proteins, cells, and viruses to chemical compounds. However, unlike antibodies, the production of aptamers is synthetic, leading to reduced batch-to-batch variability and increased cost-effectiveness[6-9]. The stability, flexibility, and synthetic nature of aptamers give them a distinct advantage in translational research, where rapid response to emerging threats is crucial.

Historically, aptamers have played a significant role in targeting medically relevant viruses. A plethora of aptamers have been generated against critical proteins and enzymes of the HIV virus, leading to advancements in both therapeutic interventions and diagnostic applications. Moreover, aptamers have been developed against a broad spectrum of viruses, including hepatitis C virus (HCV), hepatitis B virus (HBsAg), herpes simplex virus type 1 (HSV-1), human papillomavirus type 16 (HPV16), and even the Ebola virus. Recent outbreaks, such as that of the monkeypox virus, emphasize the importance of having these aptameric tools in our arsenal. The versatility of aptamers extends to respiratory viruses as well, with research showcasing their efficacy against strains like H3N2, H5N1, and H9N2 of the influenza virus, respiratory syncytial virus (RSV), and the predecessors of SARS-CoV-2, SARS-CoV, and MERS-CoV.

Our work is centered on the development of a spike-targeted aptamer against SARS-CoV-2. This aptamer has showcased promising binding efficiency with the spike protein of the virus and has demonstrated an ability to inhibit neutralization in vitro. Through the utilization of the K18-hACE2 in vivo model, we further validate the prophylaxis effect of this aptamer. As the global scientific community grapples with SARS-CoV-2 and its evolving variants, tools like aptamers stand at the forefront, promising a novel approach to diagnosis, therapy, and, ultimately, a better understanding of the virus and its pathogenicity.

## Results

### Enrichment and Identification of Aptamers Against RBD of SARS-Co-V2 Spike

Because of the variability in glycosylation modifications in the RBD of spike proteins derived from different species, which may not consistently match the SARS-CoV-2 spike protein RBD, we opted for SARS-CoV-2 spike protein RBD proteins expressed by both baculovirus-insect cells and HEK293 cells. This choice was motivated by the aim to ensure that the aptamers we obtained would be generalized effectively across varied expressions of the RBD. Our experimental process was centered around the utilization of the in vitro SELEX procedure to select 2′-Fluoropyrimidine–modified RNA aptamers that exhibited specific binding to the recombinant Spike proteins. We expressed the RBD domain of the spike protein, which included a six-histidine (His^6^) tag, in HEK293 cells. This His^6^ tag allowed for immobilization to beads, which was a pivotal step in our procedure. To first eliminate non-specific binders, we incubated the RNA aptamer library pool with agarose beads. Afterwards, supernatant was incubated with the His^6^-spike target protein to conduct a positive selection. We used PCR and in vitro transcription to amplify aptamers that successfully bound to the spike protein (Figure 1A). We conducted five rounds of selection specifically against the RBD domain. The progression of enrichment during these rounds was monitored using PCR (Figure 1B) and by detecting the bound aptamer pool to RBD (Figure 1C and D). A discernible upward trend in library binding against the RBD target was observed as we progressed through the selection rounds. Remarkably, by rounds 4 and 5, it became evident that our RNA library was enriched successfully against RBD. Given this success, we selected the 4th and 5th pools to undergo high-throughput sequencing. Our analysis of the sequencing data revealed multiple sequences showing enrichment. Of these enriched sequences, we zeroed in on 11 candidate sequences that not only displayed the highest abundance but also exhibited an increased frequency between the 4th and 5th selection rounds. The details of these sequences, along with their comparative data, are presented in Table 1 and Figure 1 E and F.

**Figure 1.**
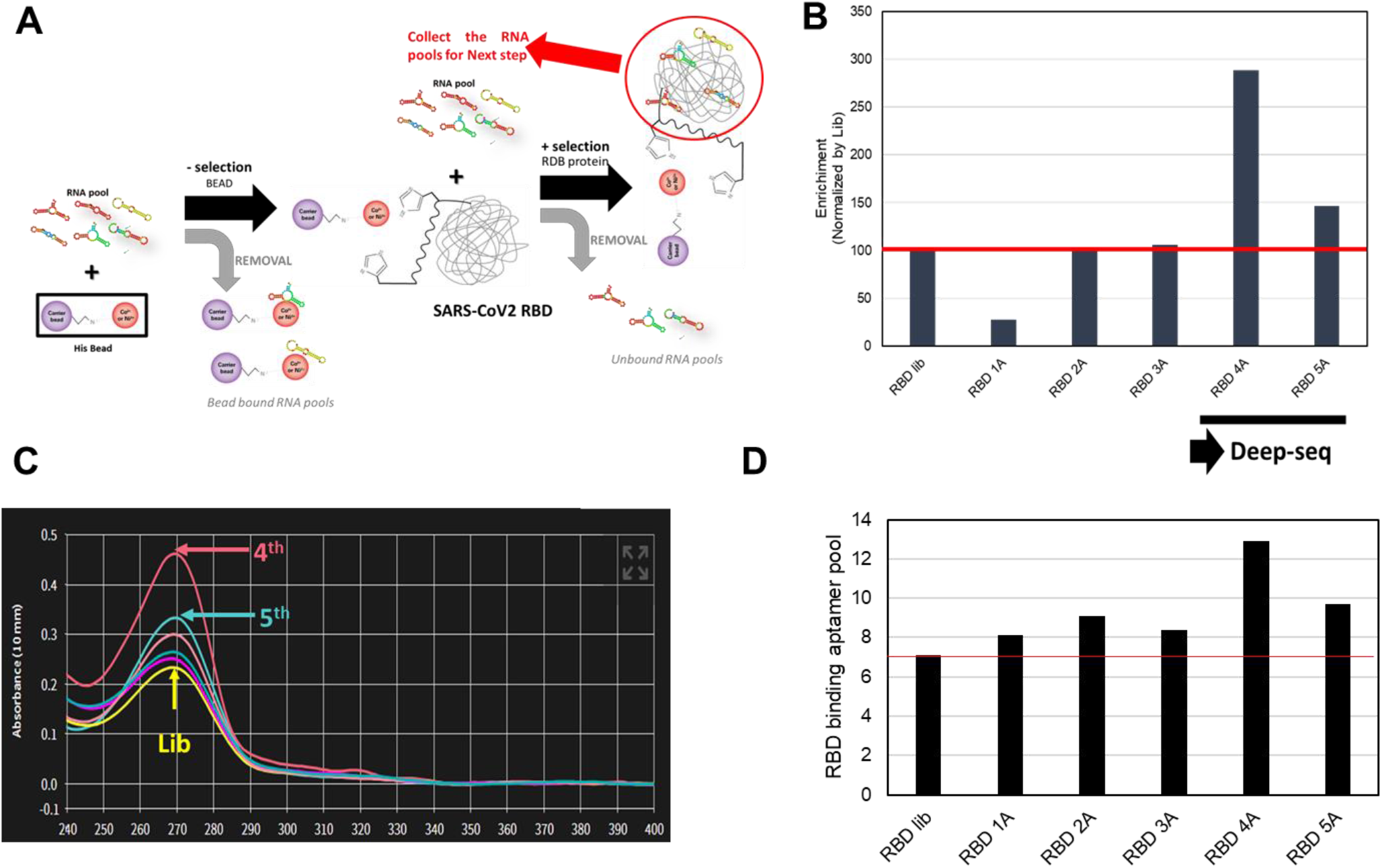
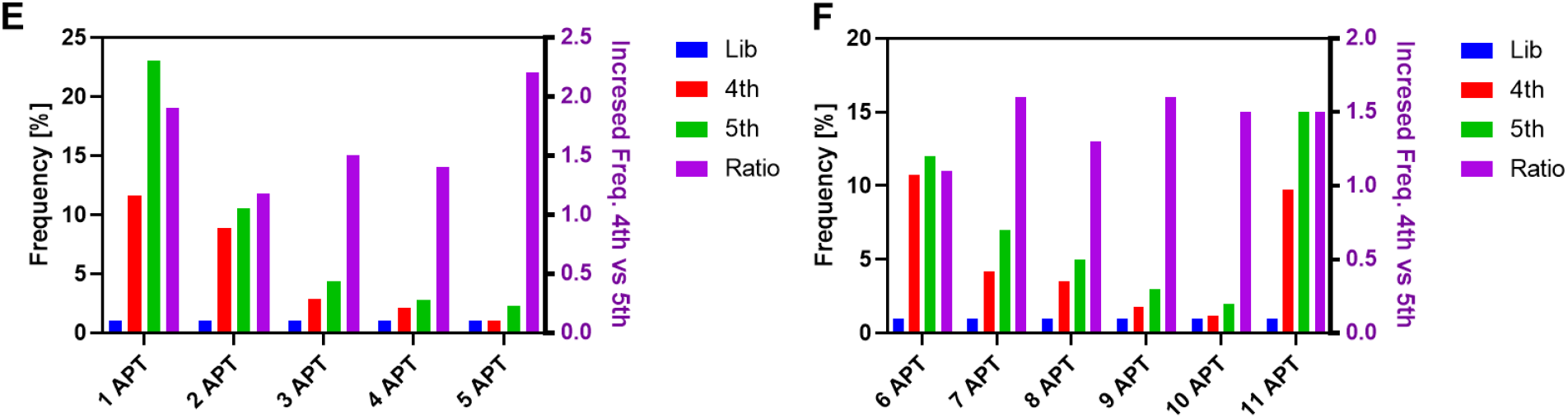
Aptamer identification and characterization. A) Protein SELEX method. B) PCR to monitor the binding increments of enriched polls with RBD-Ni-Beads (target beads) and Ni-Beads (control beads). C) Absorbance demonstrating the binding performance of each round of aptamer pools. Yellow peak, Library(Lib); red peak, 4^th^ round pools; blue peak, 5^th^ round pools. D) RNA amount to bind with RBD reacted with each round pool. E) Development of Spike RNA aptamers. Molecular enrichment at initial library (Lib), 4th round, and 5th round. After alignment of top 100 sequences, several groups of aptamers were identified. F) Frequency of each aptamer at each round. The percentage frequency of each group at each selection round was calculated by the reads of each group/the total reads of top 1000 unique sequences.

**Table 1.**
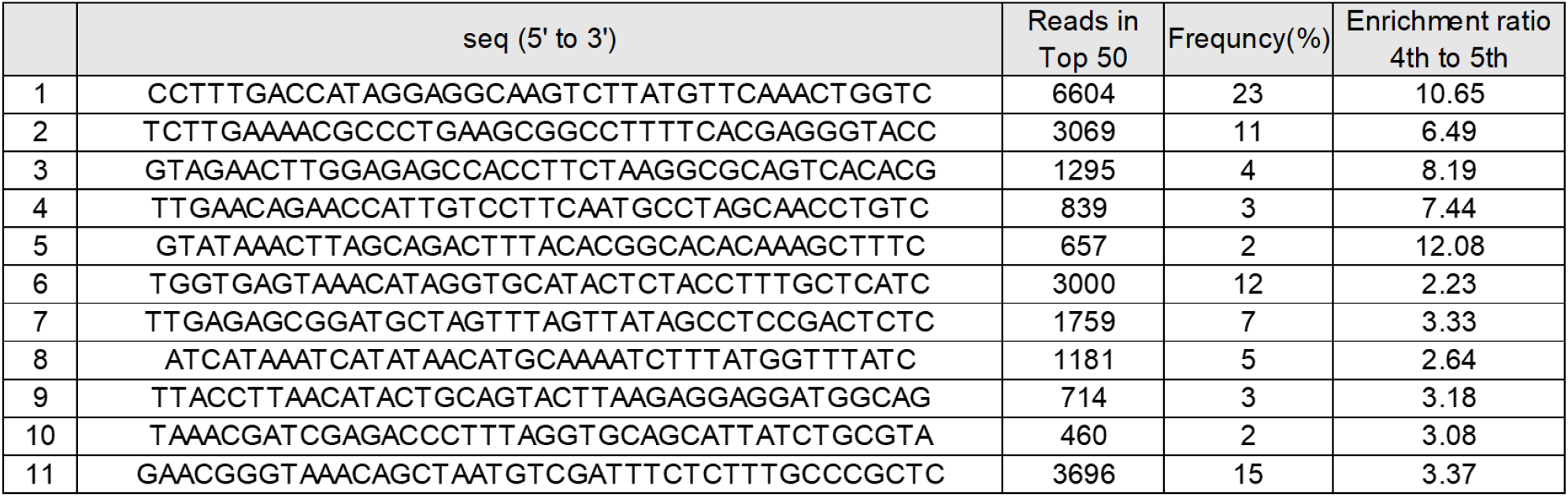
Clustering analysis of RNA libraries derived from company A was used to identify related sequence groups. After alignment of the top 100 unique sequences, several sequence groups were identified. The representative sequences of each group and their reads and frequencies are listed. The 40-nt random sequences of the RNA core regions (5′-3′) are indicated.

### Binding Affinity and Neutralization Potential of Candidate Aptamers

To ascertain the binding affinity of our candidate aptamers, we synthesized and labeled them with Cy3 fluorescent dye. These labeled aptamers were individually incubated with cells expressing either the original spike protein or the omicron variant. Subsequent binding interactions were quantified using flow cytometry. Remarkably, aptamer 1 showcased the highest binding affinity to both cells expressing the original spike protein and those expressing the omicron variant (Figure 2 A and B).

**Figure 2.**
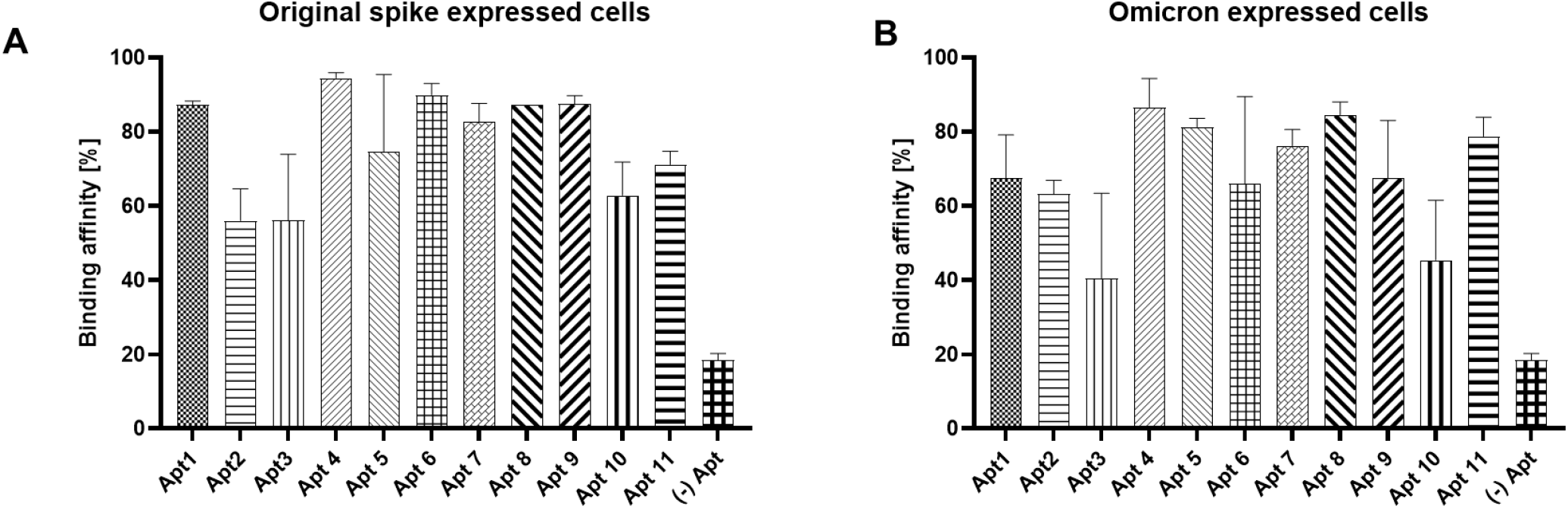
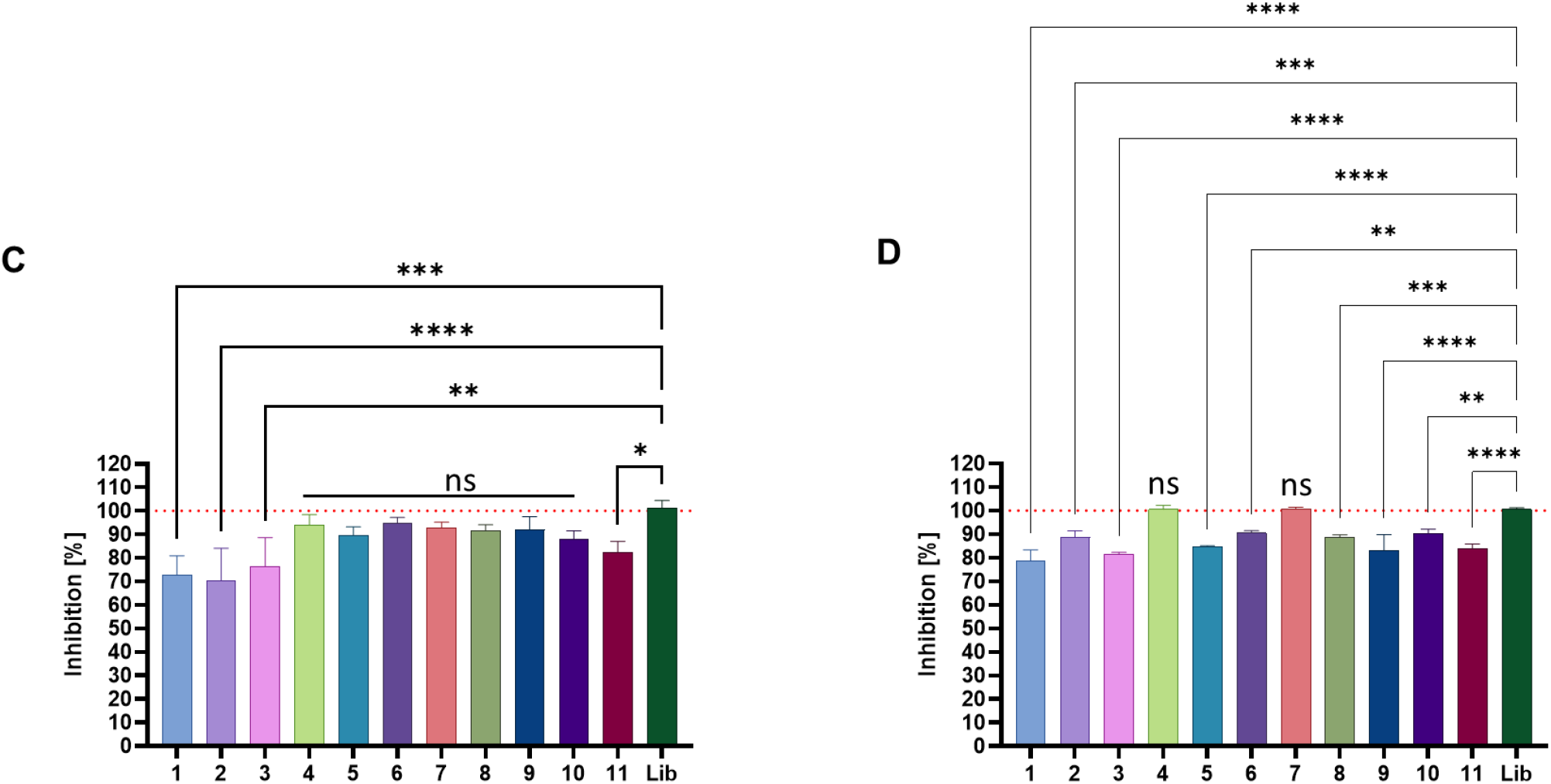
Binding ability of selected RNA aptamer clones to Spike (A) or Omicron (B) expressed HEK293 cells. (-) aptamer of targeting CD8 or each RNA aptamer clone was incubated in Spike (A) or Omicron (B) expressed HEK293 cells. The amount of RNA bound to the cells was determined by flow cytometry and represented as percentage of input RNA. (C–D) Neutralization assay with candidate aptamers. The percentage of neutralization for each tested sample in original spike (C) and omicron spike (D) expressed cells was calculated based on library-treated samples, where the library was made of 40-nt randomized RNA sequences. Data represents the mean ± standard deviation (SD). Student’s t-test, non-parametric statistical tests and ANOVA (one-way and two-way) were performed for statistical analyses. ns > 0.05, *p < 0.05, **p < 0.01, ***p < 0.001, ****p<0.0001.

To further investigate and validate the inhibitory and neutralization capabilities of the candidate aptamers, we utilized a cell fusion system. First, we generated the 293-hMyD88-Spike donor cells by transiently transfecting 293-hMyD88 cells with a spike expression plasmid that included both the wild-type spike and its variants. Then, we co-cultured these donor cells with HEK-Blue-hACE2 acceptor cells.

Notably, HEK-Blue-hACE2 cells are engineered to stably express the human ACE2 receptor, making them prone to infection. When these acceptor cells are co-cultured with the 293-hMyD88-Spike cells, the spike protein present on the donor cells binds to the ACE2 on the acceptor cells, resulting in cell fusion. Following this, we assessed the release of SEAP mediated by this fusion. HEK-Blue-ACE2 cells possess an inherent NF-κB-dependent SEAP reporter. As a result, when cell fusion occurs, MyD88 from 293-hMYD88-Spike donor cells triggers a signaling pathway in the HEK-Blue-ACE2 cells. This pathway leads to the production of SEAP in an NF-κB-dependent manner, which we subsequently measured in the co-culture supernatant using QUANTI-Blue™ Solution.

To grasp a deeper understanding, we conducted a neutralization assay using candidate aptamers. This assay hinged on the transfer of the adaptor molecule, MyD88, from a donor to an acceptor cell line. The acceptor cell line expresses an NF-κB-SEAP inducible reporter gene. Subsequently, cell fusion is easily detectable in the co-culture supernatant using the SEAP detection reagent, QUANTI-Blue Solution™. For a detailed visual representation of this mechanism, see supplementary Figure 1. This assay elucidates how an aptamer binding to the spike protein can obstruct cell fusion. This blockage prevents MyD88 from getting transferred to ACE2-expressed cells, resulting in the absence of SEAP expression and allowing for the detection of the ability of an aptamer to inhibit neutralization. From our results, it became evident that introducing aptamer number 1 considerably diminished the expression of SEAP in cells expressing either the original or omicron spike protein (Figure 2C and D). This outcome highlights the potential of aptamer number 1 to be a groundbreaking neutralizing antiviral agent against SARS-CoV2 infection.

### Determining the Binding Affinity of Selected RNA Aptamers to SARS-CoV-2 Spike Proteins and its Omicron Variant

To assess the binding affinity of the identified aptamers for the spike protein and its omicron variant, representative sequences from each group were synthesized and subjected to flow cytometry analysis. Our findings demonstrated that these RNA aptamers exhibited selective binding to HEK293 cells expressing either the original spike or the omicron variant. This result underscores aptamers specificity towards the spike protein. Notably, the dissociation constant (Kd) for Apt1 was recorded as 37.71 nM for the original spike and 31.92 nM for the omicron variant (Figure 3 and Table 2).

**Figure 3.**
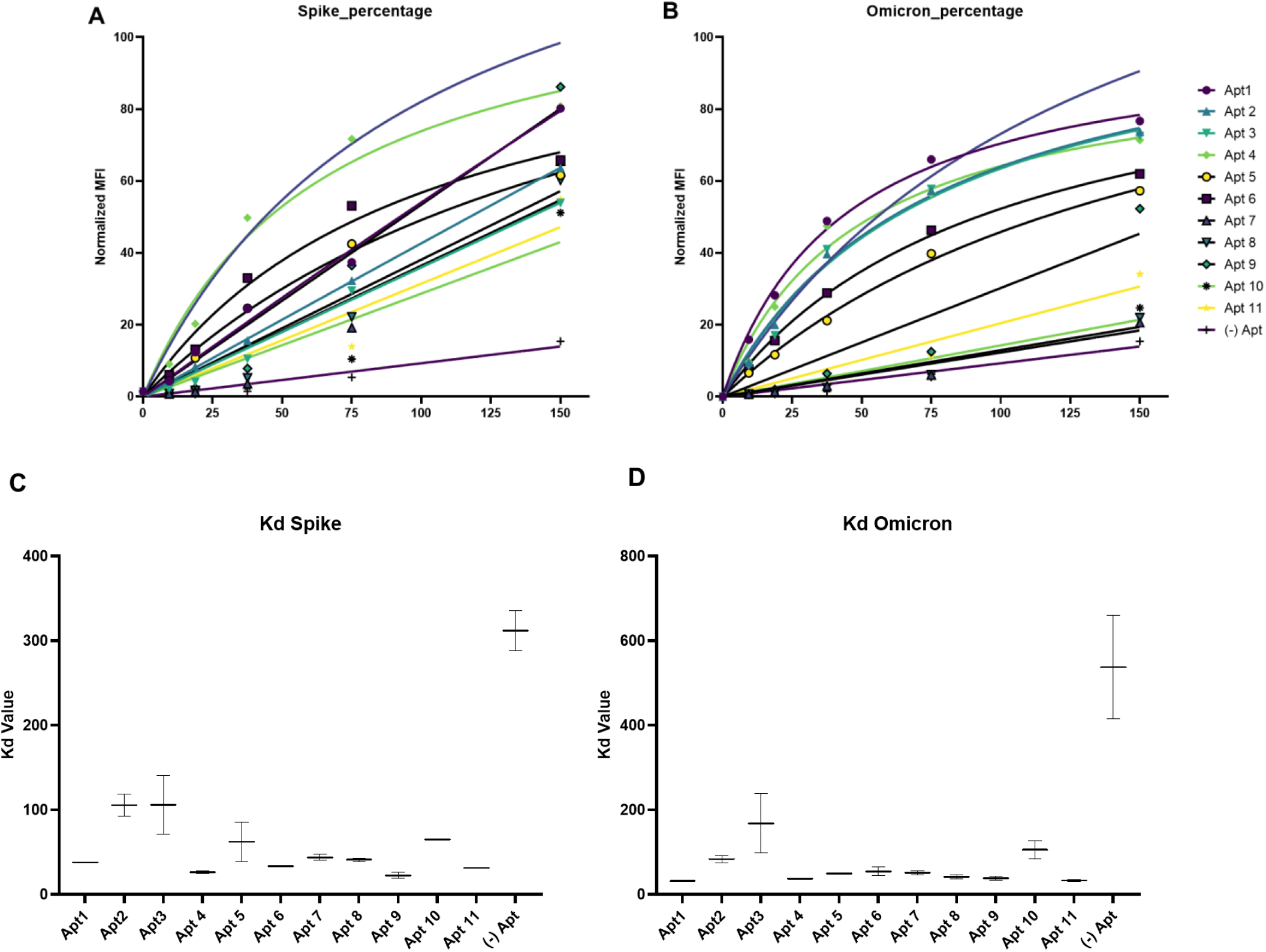
Fluorescence binding curves for each RNA aptamer determined by titrating RNA aptamer concentration. Binding curves of Cy3-labeled aptamers on Spike (A) or Omicron (B) expressed HEK293 cells for calculation of the apparent Kd of aptamer-cell interaction. Binding was analyzed using flow cytometric assay. (C–D) Comparison of the KD values for binding of RNA aptamers to original spike or omicron expressed HEK293 cells. The KD values are the mean ± SD of 2–3 independent experiments.

**Table 2.**
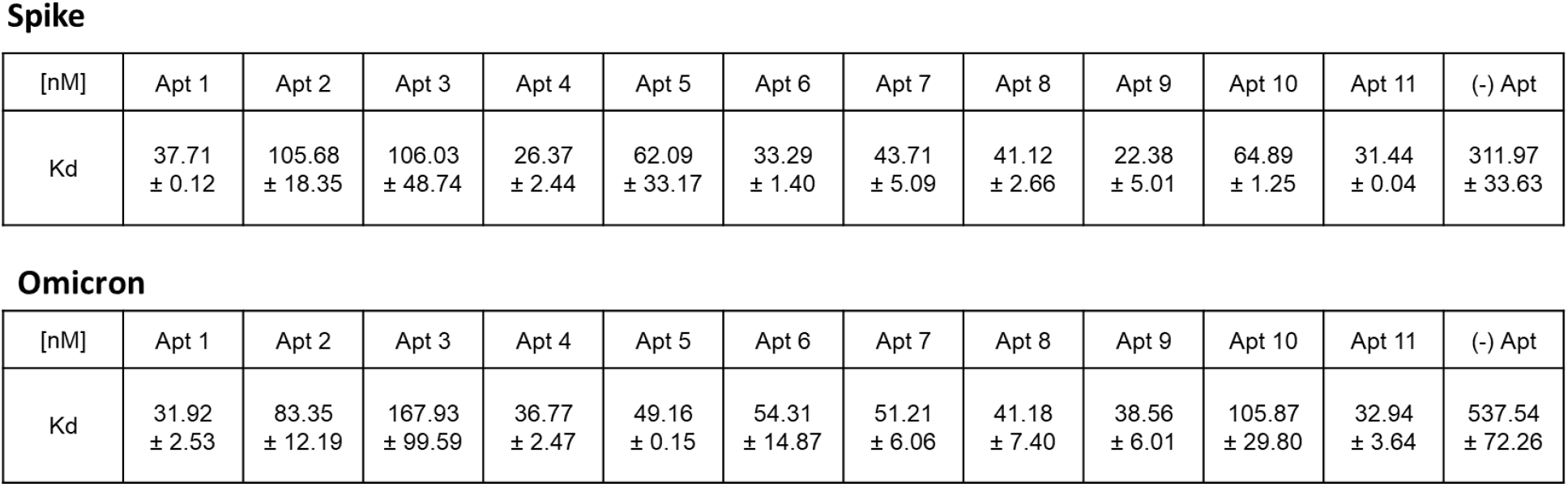
Michaelis-Menten binding curves to evaluate Kd (nM) were performed. Data shown are mean ± SD of three independent experiments.

Spike
| [nM] | Apt 1 | Apt 2 | Apt 3 | Apt 4 | Apt 5 | Apt 6 | Apt 7 | Apt 8 | Apt 9 | Apt 10 | Apt 11 | (-) Apt |
| --- | --- | --- | --- | --- | --- | --- | --- | --- | --- | --- | --- | --- |
| $K_d$ | 37.71<br>± 0.12 | 105.68<br>± 18.35 | 106.03<br>± 48.74 | 26.37<br>± 2.44 | 62.09<br>± 33.17 | 33.29<br>± 1.40 | 43.71<br>± 5.09 | 41.12<br>± 2.66 | 22.38<br>± 5.01 | 64.89<br>± 1.25 | 31.44<br>± 0.04 | 311.97<br>± 33.63 |

**Table 2.** Michaelis-Menten binding curves to evaluate $K_d$ (nM) were performed. Data shown are mean ± SD of three independent experiments.
| [nM] | Apt 1 | Apt 2 | Apt 3 | Apt 4 | Apt 5 | Apt 6 | Apt 7 | Apt 8 | Apt 9 | Apt 10 | Apt 11 | (-) Apt |
| --- | --- | --- | --- | --- | --- | --- | --- | --- | --- | --- | --- | --- |
| $K_d$ | 31.92<br>± 2.53 | 83.35<br>± 12.19 | 167.93<br>± 99.59 | 36.77<br>± 2.47 | 49.16<br>± 0.15 | 54.31<br>± 14.87 | 51.21<br>± 6.06 | 41.18<br>± 7.40 | 38.56<br>± 6.01 | 105.87<br>± 29.80 | 32.94<br>± 3.64 | 537.54<br>± 72.26 |

### Development of truncated aptamers

### In vitro neutralization by Apt1 and tApt1

### Enhanced Binding and Internalization of Truncated Apt1 (tApt1) Aptamers to Original and Omicron Spike

Using computational prediction based on structural analysis, we streamlined the Apt1 aptamer to its smallest functional unit while ensuring it retained its binding affinity to the spike protein. We derived four truncated versions of Apt1, referred to as tApt1, each originating from complementary sequences of the 72-mer Apt1 aptamer (Figure 4). When assessing affinity, the tApt1 showed a superior binding capacity compared to full-length Apt1, with Kd values of 20.73 nM for the original spike and 20.80 nM for the omicron spike variant (Figure 4B). This enhanced affinity with tApt1 suggests that the bases we eliminated from the parent aptamer’s terminal ends were superfluous. More crucially, their removal did not compromise the functional structure essential for aptamer binding and recognition. Additionally, tApt1 demonstrated comparable neutralization inhibition as its predecessor in the cell fusion system (Figure 4C). Given our goal to utilize an anti-spike aptamer therapeutically, we conducted cell internalization assays to assess the potential of tApt1 in targeted delivery. We tested the uptake of Cy3-labeled Apt1 and tApt1 aptamers by incubating them at a concentration of 200 nM with spike-expressing HEK293 cells. As a control, we treated cells with a Cy3-labeled CD8 aptamer in a manner similar to the flow cytometry tests. One hour post-incubation, we detected distinct punctate fluorescence patterns in both the cytoplasm and nuclei of cells, indicating successful internalization of Apt1 and tApt1 into spike-expressing cells (Supplementary Figure 2). These observations highlight the improved binding and functionality of tApt1 for both the original and omicron spike variants.

**Figure 4.**
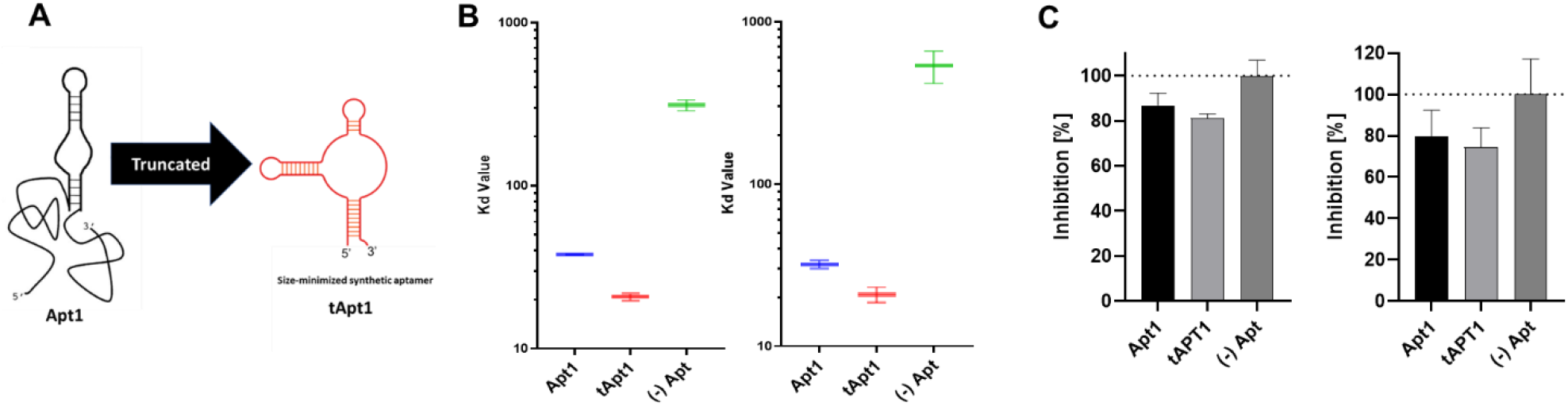
Schematic generation of truncated aptamer (tApt1) and tApt1 characterization.

### Inhalation of tApt1 effectively treats SARS-CoV-2 infection in hACE2 mice

To assess the therapeutic efficacy of inhaled tApt1 in vivo, we employed hACE2-transgenic mouse models. Prior to SARS-CoV-2 inoculation, mice were administered with tApt1 through inhalation at time points of -6, -3, and 0 days, as depicted in Figure 5A. Administration was facilitated using a specialized mouse inhalation system, represented in Figure 5B. Notably, following SARS-CoV-2 challenge, tApt1-inhaled mice exhibited resistance to weight loss on both day 1 (Figure 5C) and day 5 (Figure 5D), in contrast to their counterparts treated with a negative control aptamer.

**Figure 5.**
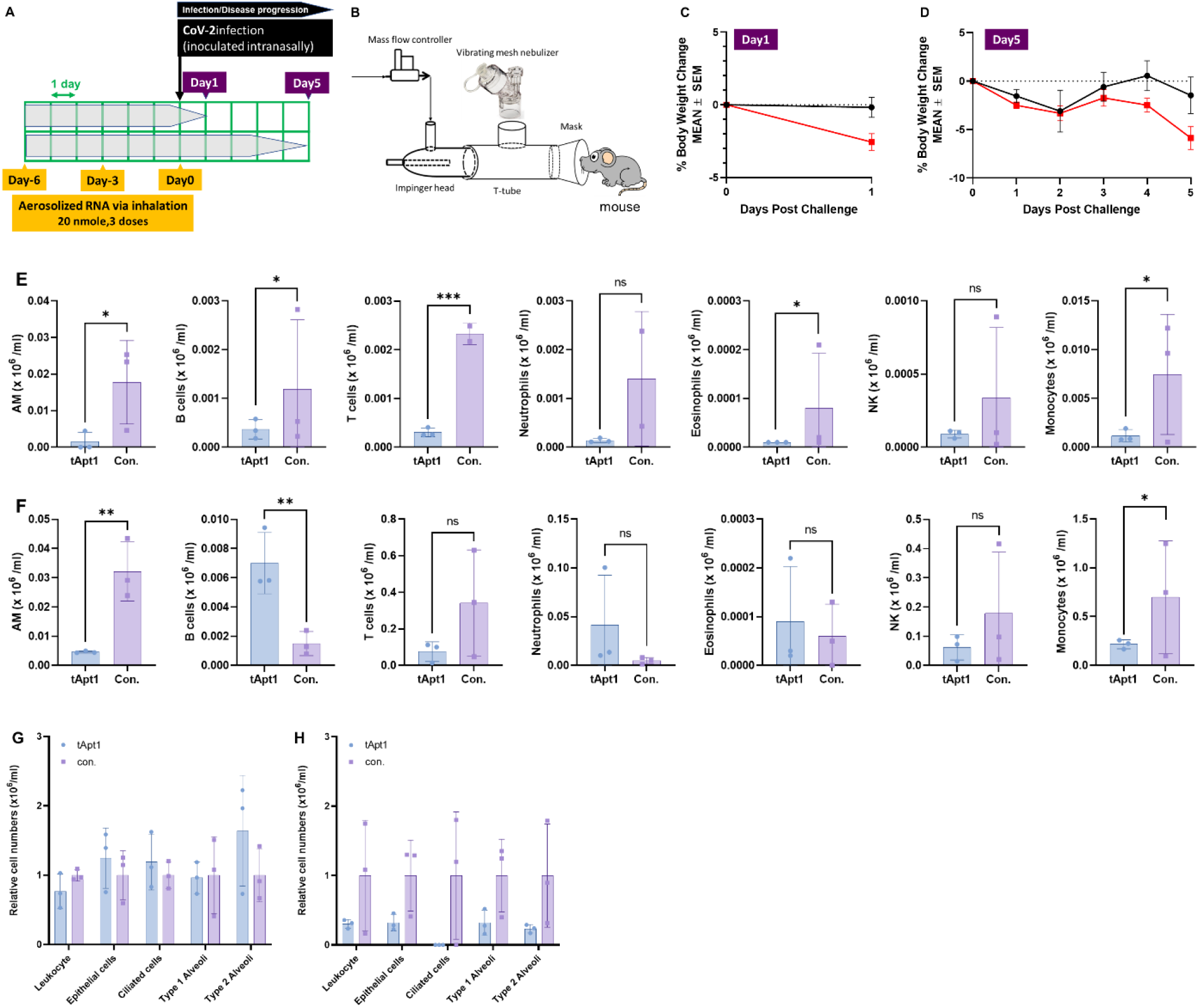
Inhalation administration of tApt1 into K18-hACE2 mice. A) schematic of inhalation of a multidose regimen of aerosolized RNAs.

An overactive pro-inflammatory response to SARS-CoV-2 infection might be the underlying cause for the pulmonary complications and subsequent respiratory distress seen in some COVID-19 patients [12]. Notably, after recovering from COVID-19, an elevated cell count has been observed in patients’ airways, largely due to an increased presence of airway macrophages (AMs), T cells, and B cells [13].To gain insight into the immune cell landscape following SARS-CoV-2 infection in K18-hACE2 mice, we undertook flow cytometric analysis of bronchoalveolar lavage (BAL) fluid, sampled at two distinct post-inoculation intervals (Figures 5E and F). Encouragingly, mice administered with inhaled tApt1 showcased substantial suppression across various cell types, including leukocytes, endothelial cells, epithelial cells, ciliated epithelial cells, and both type I and II alveoli on days 1 and 5 (Figures 5G and H). Moreover, a striking outcome was the absence of live virus in the brain and pancreas of tApt1-inhaled mice (Figure 6), underscoring the potential of inhaled tApt1 in thwarting SARS-CoV2 infection. Collectively, our results illuminate the potential therapeutic role of inhaled tApt1 against SARS-CoV2 infection. The benefits span protection against weight loss, diminished viral load, and modulation of the immune response elicited by the virus.

**Figure 6.**
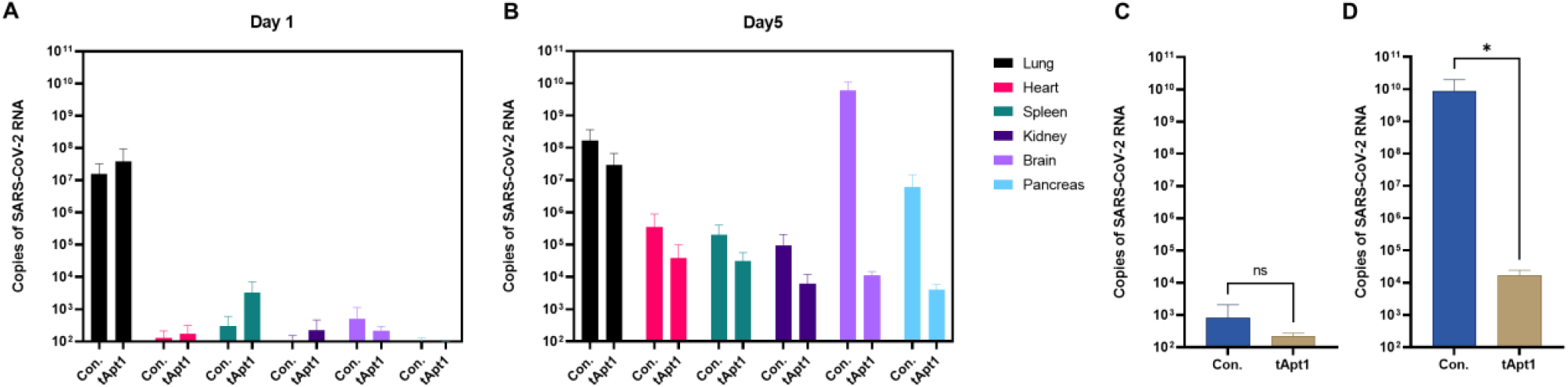
Detection of viral mRNA in mouse tissue.

## Discussion

Recent studies have delved into various therapeutic strategies targeting SARS-CoV-2 infection, with a particular emphasis on virus-cell interaction dynamics. A notable approach involves the use of aptamers that specifically bind to the SARS-CoV-2 spike protein. By doing so, these aptamers impede the interaction between the ACE2 receptor on human cells and the spike protein, thereby curtailing the virus’s ability to infect host cells. Understanding the transmission dynamics of SARS-CoV-2 is crucial for devising effective preventive strategies. A dominant route of transmission is through respiratory droplets, particularly sneezes. Remarkably, a single sneeze can release up to 10,000 droplets, presenting significant potential for viral spread, given that the virus predominantly resides in the airways (Figure 7). While our current focus is on SARS-CoV-2, it’s essential to look beyond to other viruses. The World Health Organization (WHO) has recently emphasized the need to prepare for potential pandemics caused by the pan-sarbecovirus group. Viruses from this group have an inherent propensity to spread via the respiratory system, primarily infecting individuals through inhalation. The inhaled virus can then readily attach to and colonize the airways, specifically the nose and lungs. The concept of inhaled aptamers showcases promise as they can potentially obstruct viral receptors, preventing them from engaging with human cell receptors and thus inhibiting infection.

**Figure 7.**
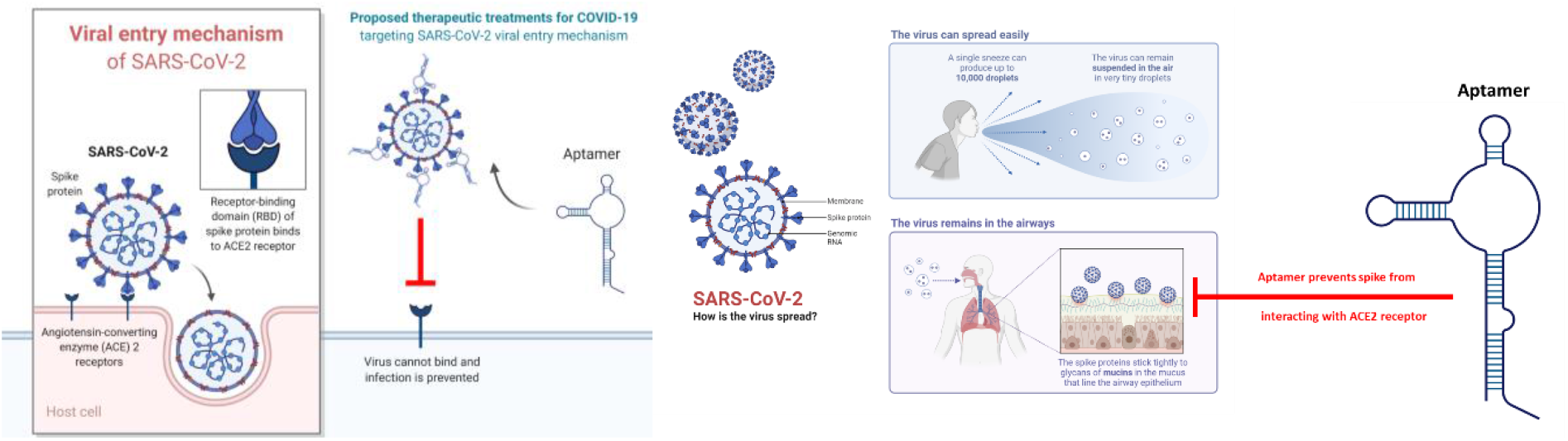
Overview of therapeutic strategies to treat SARS-CoV-2 infection based on virus-cell interactions. The aptamer bound to SARS-CoV-2 spike inhibits the interaction between ACE2 and Spike and thus prevents infection. SARS-CoV2 can spread easily by a sneeze. A single sneeze can produce up to 10 thousand droplets. The virus remains in the airways. It does not belong to SARS-CoV2. Recently the WHO announced we need to prepare the NEXT pandemic for pan-sarbecovirus. This type of virus easily spread to the airways. You breathe air and virus spread into your nose and lung. Inhaled aptamer can prevents viral receptor from interacting with human receptor.

Inhalation drug delivery offers several advantages that enhance therapeutic outcomes. A foremost benefit is its non-invasive, needle-free nature, which provides an alternative to traditional injections, alleviating the associated discomfort and potential complications (Supplementary Figure 3). Such a mode of delivery undoubtedly enhances the patient experience. Additionally, the lungs, with their vast surface area and permeable epithelium, serve as an efficient gateway for inhaled therapeutics. The direct administration of drugs to the lungs ensures targeted delivery to the site of action. This not only accelerates the onset of therapeutic effects but also allows for the use of smaller drug dosages, achieving optimal drug concentrations right where needed. Another pivotal advantage of inhalation delivery is its ability to minimize systemic bioavailability. This reduction mitigates the risk of systemic toxicities. Furthermore, the inhalation route circumvents first-pass metabolism in the liver, a process that can compromise drug efficacy and necessitate increased dosages in other modes of administration.

## Supplementary Figures

**Supplementary Figure 1.**
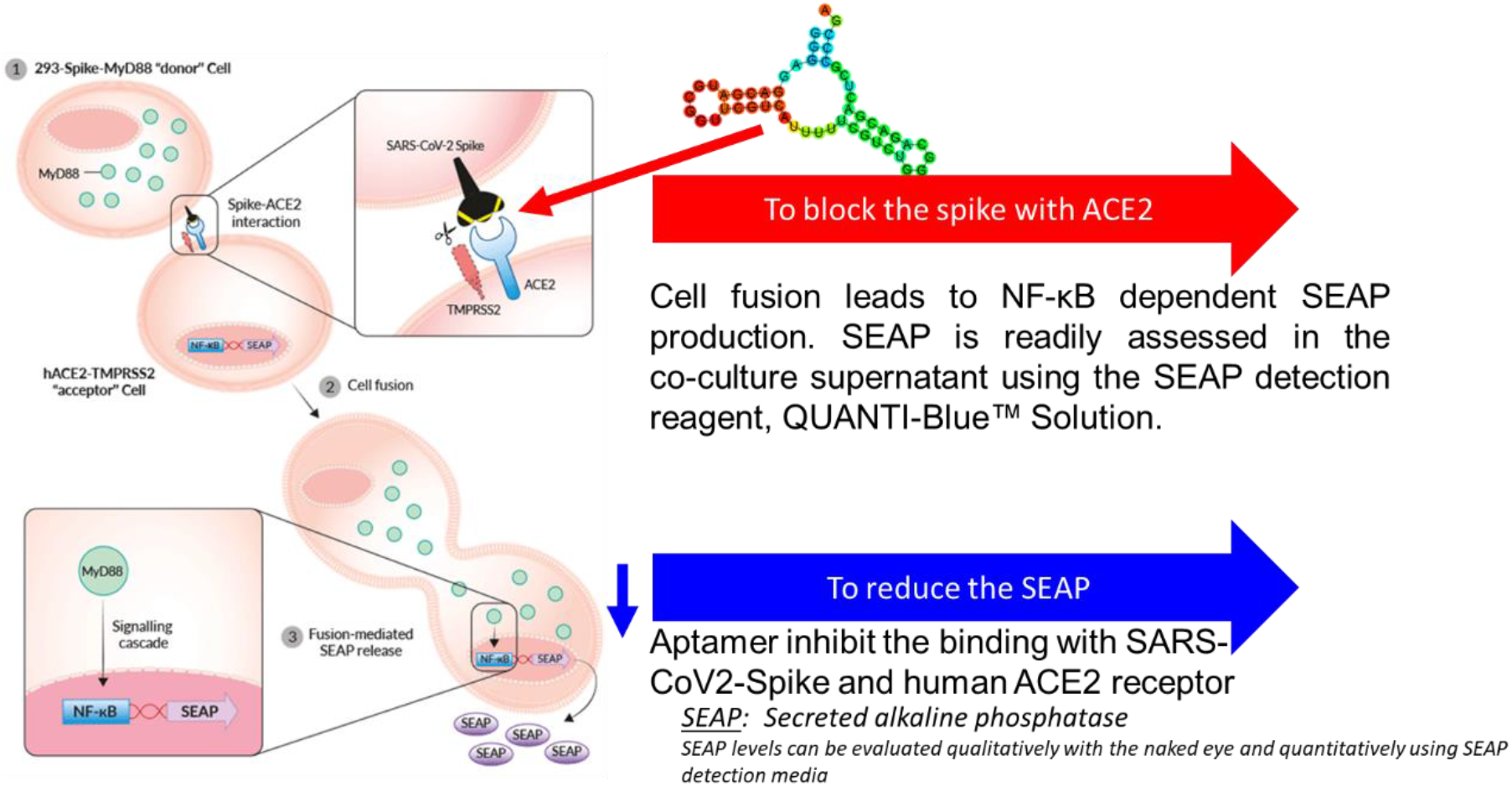
Neutralization assay with candidate aptamers. This assay relies on the transfer of the adaptor molecule, MyD88, from a donor cell line to an acceptor cell line expressing an NF-kB -SEAP inducible reporter gene. Cell fusion is then readily assessable in co-culture supernatant using the SEAP detection reagent, QUANTI-Blue Solution™. Left side from Invivogen webpage. The figure looks at this method more closely. Aptamer binds with spike protein and block the cell fusion. This means myD88 doesn’t transfer into ACE2 expressed cells and SEAP doesn’t express. We can easily detect the Aptamer function to inhibit neutralization.

**Supplementary Figure 2.**
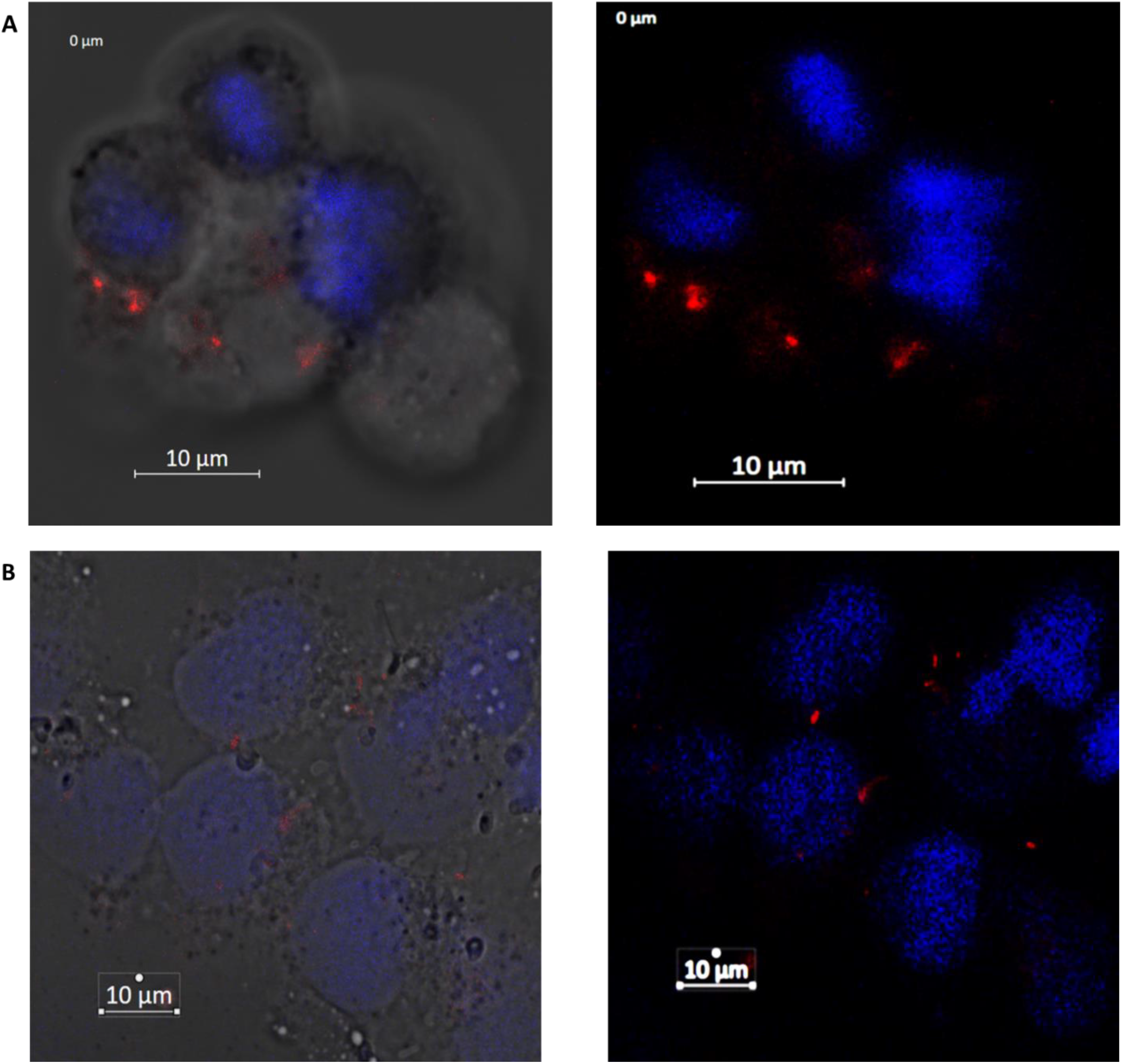
Cell binding of APT1(A) and tAPT1(B) aptamers. Representative confocal images of spike expressed HEK293 cells incubated for 60 min in a cell culture incubator with Cye-labeled APT1. After washing and fixation, cells were labeled with NucBlue™ Live ReadyProbes™ Reagent to visualize the cell membrane, and with DAPI (blue) to stain nuclei. Cy3-labeled aptamers are displayed in red. All digital images were captured at the same setting to allow direct comparison of staining patterns. Magnification 63×, 1.0× digital zoom, scale bar = 10 μm. At least three independent experiments were performed.

**Supplementary Figure 3.**
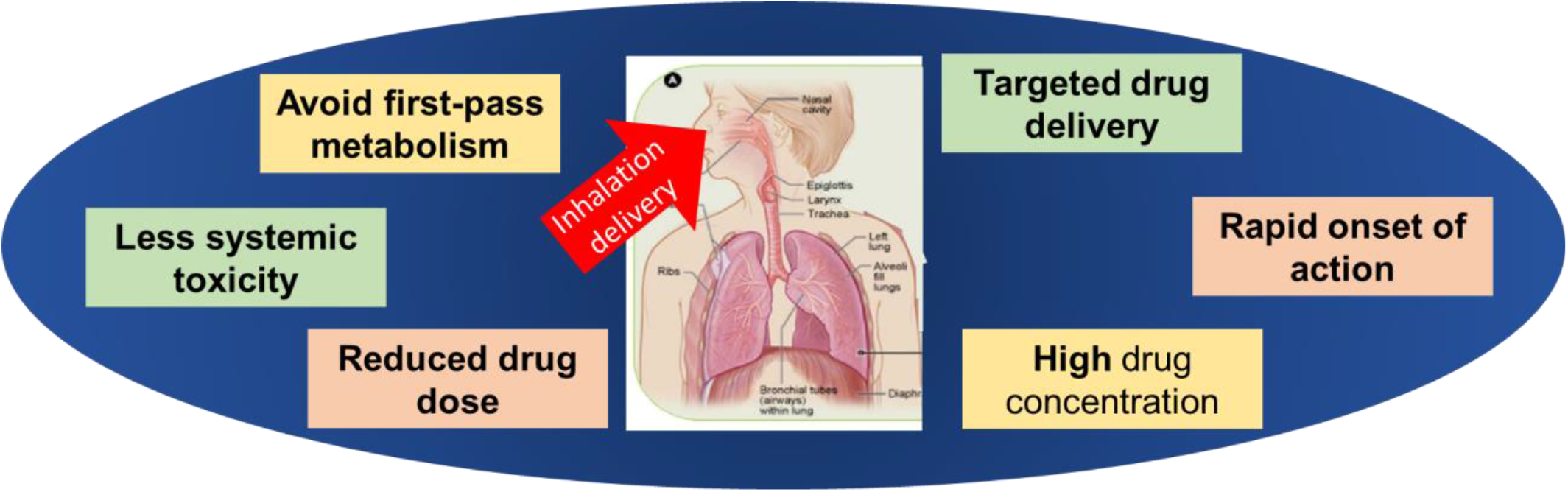
Inhalation delivery offers several distinct advantages over other routes of administration.

